# Locus-associated R-loop repression in *cis*

**DOI:** 10.1101/2021.05.25.445644

**Authors:** Negin Khosraviani, Karan J. Abraham, Janet N.Y. Chan, Karim Mekhail

## Abstract

R-loops exert varied beneficial or detrimental effects. To assess the function of an R-loop at a specific genetic locus, we had developed an inducible RNaseH1-EGFP-dCas9 (RED) protein chimaera as part of a locus-associated R-loop-modulating system (LasR). LasR is compatible with R-loop modulation in *trans*, which targets RED to one locus to repress R-loops at another spatially proximal site. Here we use the LasR system for R-loop modulation in *cis*, which consists of targeting RED directly to an R-loop. The combination of LasR in *cis* and *trans* will be essential to ascribe functions to specific R-loops within varied molecular contexts and study designs.

## Text

R-loops are three stranded nucleic acid structures harboring an RNA-DNA hybrid and a displaced single-stranded DNA^1,2^. R-loops are common by-products of transcription and their levels can exert varied beneficial or harmful effects depending on the specific molecular context^1,2^. Most of our current understanding of R-loop functions stems from experimental approaches that evaluate phenotypic changes following the repression of R-loops via the overexpression of the R-loop repressor ribonuclease H1 (RNase H1). However, such an approach has a major limitation-observed phenotypic changes may be the result of the repression of R-loops at the locus of interest or at other R-loop-containing loci elsewhere in the genome. To overcome this limitation, we previously developed a tetracycline-inducible RNaseH1-EGFP-dCas9 (RED) protein chimaera as part of a locus-associated R-loop modulation system (dubbed RED-LasR or LasR)^3^. In this system, the RED fusion protein can be expressed at the minimum necessary level and enriched at the locus of interest using short guide RNA (sgRNA)^3-5^. As controls, we use a version of RED called dRED, in which the RNaseH1 moiety is catalytically dead with a D210N point mutation (dRNaseH1)^3,6^. The inactivated enzyme loses the hybrid-suppressing ability and in some cases even results in increased R-loop levels, likely by acting as a dominant-negative that binds R-loops preventing its removal by endogenous R-loop repressors^1,3,7,8^. Other negative controls include using non-targeting sgRNA controls (sgNT) and assessing R-loop levels at non-targeted or unrelated DNA loci^3^.

We previously used the LasR system to achieve R-loop modulation in *trans* by taking advantage of extensively characterized human chromosome loops^3^. Specifically, targeting RED or dRED to the intergenic spacer site 28 kb from the precursor-rRNA transcription start site of ribosomal DNA repeats (IGS28) repressed and increased R-loop levels at the spatially proximal IGS18 site, respectively, without impeding local transcription. Using this approach, we discovered that IGS18 R-loops are deposited by nucleolar RNA Polymerase II to prevent RNA Polymerase I from synthesizing sense intergenic non-coding RNAs (sincRNA) that compromise the ribosome biogenesis-conducive liquid-liquid phase separation properties of the nucleolus^3^. Use of LasR to achieve R-loop modulation in *cis* would further broaden the usability and accessibility of this tool.

To assess whether the LasR system can be used to achieve R-loop modulation in *cis*, we first assessed the targeting of RED or dRED to the extensively characterized R-loop at the 5’Pause site of the *β-ACTIN* locus^9,10^. Using our previously reported^3^ tetracycline concentration of 1 μg/mL and a pool of sgRNAs targeting both DNA strands of this R-loop (sgACTIN), chromatin immunoprecipitation (ChIP) showed a similar enrichment of RED and dRED at the targeted site compared to sgNT (Fig. 1a,b). Targeting RED to the site (Fig. 1a) significantly decreased its R-loop levels (−42%, Fig. 1c) as revealed by DNA-RNA immunoprecipitation (DRIP). In contrast, targeting dRED to the site (Fig. 1b) only qualitatively increased its R-loop levels (+75%, Fig. 1d). The significant RED-dependent decrease in R-loop levels at the site was not accompanied by changes in transcript levels (Fig. 1e). Also, targeting RED to the site with sgACTIN did not alter R-loop levels at the unrelated R-loop-containing PMS2-TSS, RPL13A, and LINE1 sites tested (Fig. 1f)^11-13^. Decreasing the concentration of tetracycline abolished the R-loop-modulating capacity of the LasR system (Fig. 1g,h). In contrast to the use of a pool of sgRNAs targeting both DNA strands (Fig. 1), strand-specific sgRNA pools failed to enrich RED or alter RNA and R-loop levels at the targeted site (Fig. 2). Together, these findings indicate that the LasR system can be used with a pool of sgRNAs targeting both DNA strands to achieve locus-associated modulation of R-loop levels in *cis*.

**Fig. 1.**
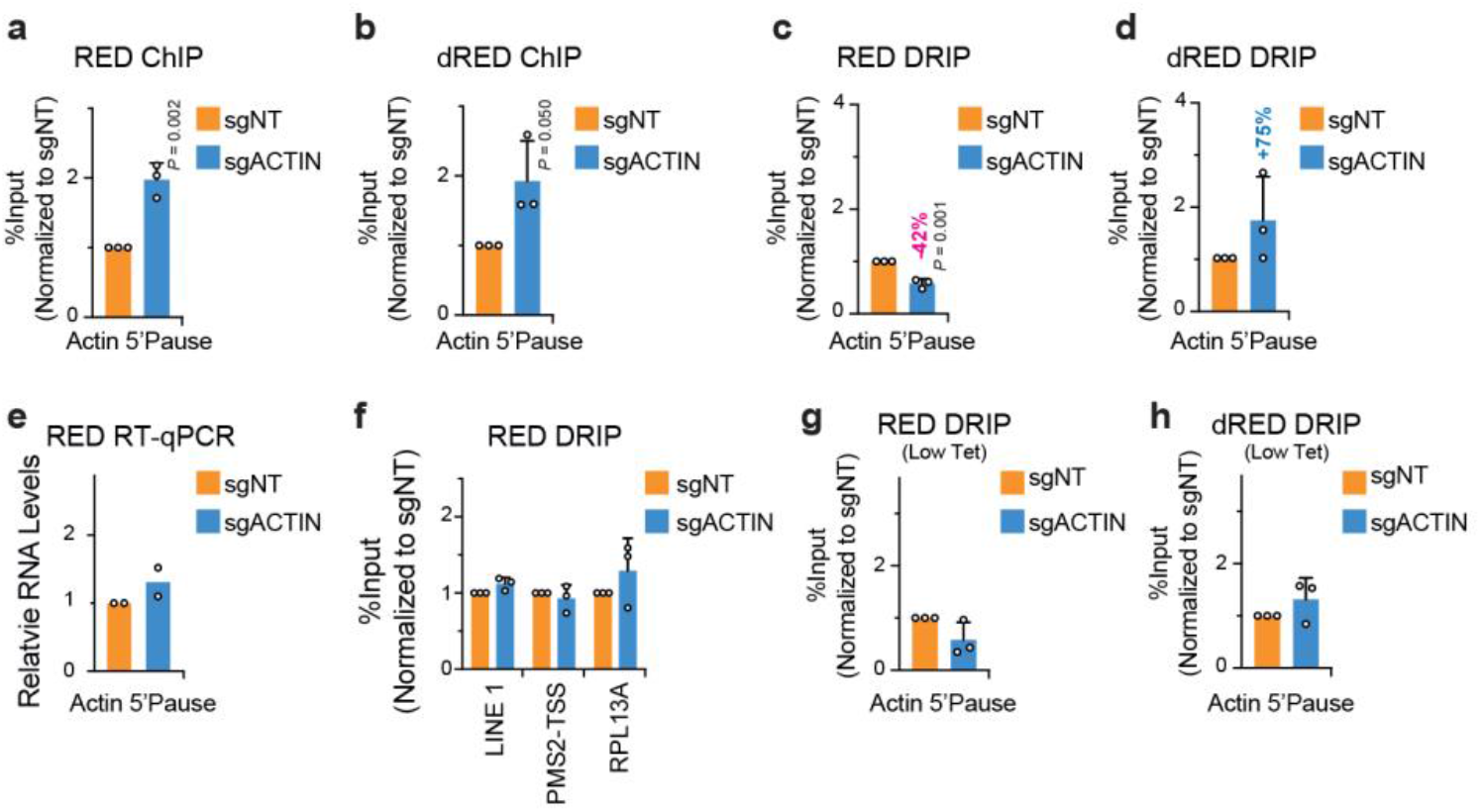
Locus-associated R-loop repression in *cis*. **a, b**, The short guide RNAs for the *β-ACTIN 5’ Pause* (sgACTIN) enriched the tetracycline (1 μg/mL Tet)-induced RED (**a**) and dRED (**b**) at the same locus in anti-GFP ChIP, using IgG as background control. Enrichments are normalized to non-targeting control signal (sgNT). **c, d**, DRIP analysis shows that using RED (**c**) or dRED (**d**) together with sgACTIN respectively decreased and increase R-loop levels in *cis*. **e**, Using RED together with sgACTIN did not alter local transcript levels. **f**, Using RED together with sgACTIN did not change R-loop levels at *LINE1, PMS2-TSS*, or *RPL13A*. **g, h**, Decreasing the Tet concentration abrogates the chimaera-dependent *cis* R-loop modulation by sgACTIN. **a-i**, HEK293T cells; data are shown as means ± s.d.; two-tailed *t*-test **(a-e, g, h)** and two-tailed ANOVA with Tuckey’s multiple comparisons test **(f);** n=3 (**a-d, f-h**) and n=2 (**e**) biologically independent replicates.

**Fig. 2.**
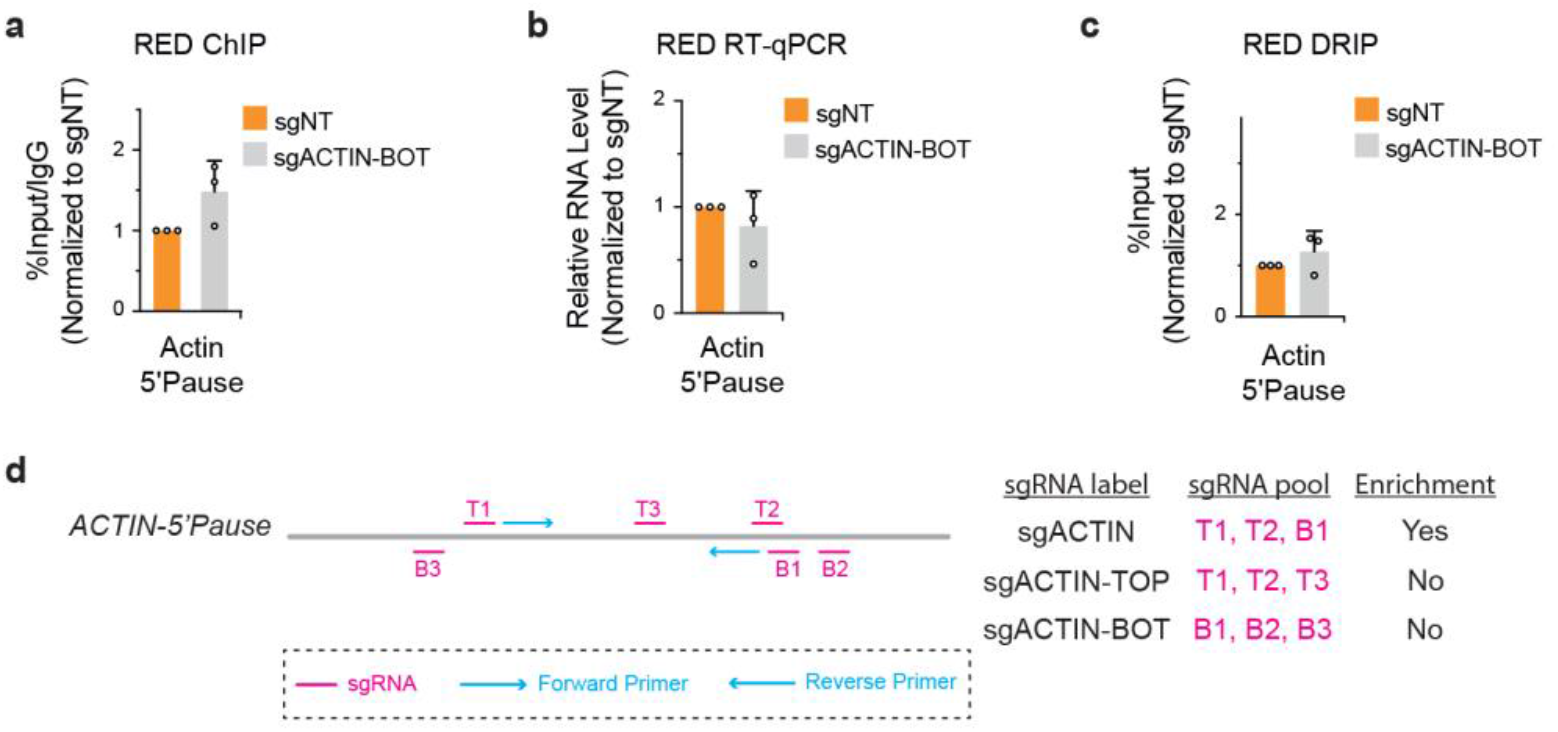
Using the LasR system with strand-specific or mixed pools of sgRNAs. **a-d**, Representative data and summary of experiments testing strand-specific short guide RNAs complementary to either strand of the *β*-ACTIN 5’Pause R-loop (sgACTIN-TOP or sgACTIN-BOT). HEK293T cells; data are shown as means ± s.d.; two-tailed *t*-test; n=3 technical replicates **(a)** and n=3 biologically independent replicates **(b**,**c). a**, Unlike sgACTIN, sgACTIN-BOT did not enrich the tetracycline (1 μg/mL Tet)-induced RED at the *β*-ACTIN 5’Pause in anti-GFP ChIP, using IgG as background control. Enrichments are normalized to non-targeting control signal (sgNT). **b, c**, Using sgACTIN-BOT together with RED did not alter the levels of transcripts (**b**) or R-loops (**c**) at the *β*-ACTIN 5’Pause. **d**, Schematic summarizing the ChIP enrichment of RED and dRED at the *β*-ACTIN 5’Pause site using the sgRNA pools of sgACTIN, sgACTIN-TOP, and sgACTIN-BOT. The positions of the sgRNAs and qPCR primers used in ChIP are illustrated.

Together with earlier work^3,14^, the findings presented here highlight the efficacy and flexibility of the LasR system. Specifically, the data show that the system is amenable to modulating the levels of R-loops associated with specific genetic loci in *cis* or *trans*. Use of the LasR system to achieve R-loop modulation in *cis* makes this tool more accessible to researchers with limited knowledge of genome folding principles and data, some of which requires an advanced knowledge of chromosome conformation capture, genome-wide sequencing, and related technologies^15^. The *cis* approach would also be necessary when the studied R-loop falls within a genetic locus with weak or no intrachromosomal interactions. On the other hand, use of the LasR system to achieve R-loop modulation in *trans* supplies a number of benefits and may be preferred depending on the exact experimental and molecular context. For instance, the *trans* approach may be preferred when targeting RED or dRED directly to an R-loop of interest either artificially compromises local transcription and other resident chromosomal features. The *trans* application may also prove essential if targeting of RED or dRED in *cis* fails due to unforeseen local steric hindrance or weakly accessible silent chromatin environments. In conclusion, the ability of the LasR system to achieve R-loop modulation in *cis* and *trans* will provide researchers with a high degree of flexibility when designing experiments requiring locus-associated modulation of R-loop levels. These approaches promise to pave the way towards the functional characterization of individual R-loops across the genome, shedding new light on human health and disease.

## Methods

### Cell culture and general materials

HEK293T T-REx™ cells (ThermoFisher Scientific) were cultured in Dulbecco’s modified Eagle medium (DMEM; Wisent Bioproducts) supplemented with 10% tetracycline-free fetal-bovine serum (FBS; Wisent) and 1% penicillin/streptomycin (Wisent). Cells were cultured at 37°C in a humidified atmosphere with 5% CO_2_.

### Transfection of RED-LasR system

HEK293T T-REx™ cells were transfected as previously described^3^. In brief, cells that were grown to 70% confluence were incubated with medium containing tetracycline (1μg/ml) and transfected with RNH1-EGFP-dCas9 (RED) or dRNH1-EGFP-dCas9 (dRED) using Lipofectamine3000 (ThermoFisher) as per the manufacturer’s instructions. Induced cells were also co-transfected with sgRNAs, listed in Table 1, using RNAiMAX (ThermoFisher) as per the manufacturer’s instructions. Cells were then incubated for 36 h before further experiments were performed.

**Table 1.**
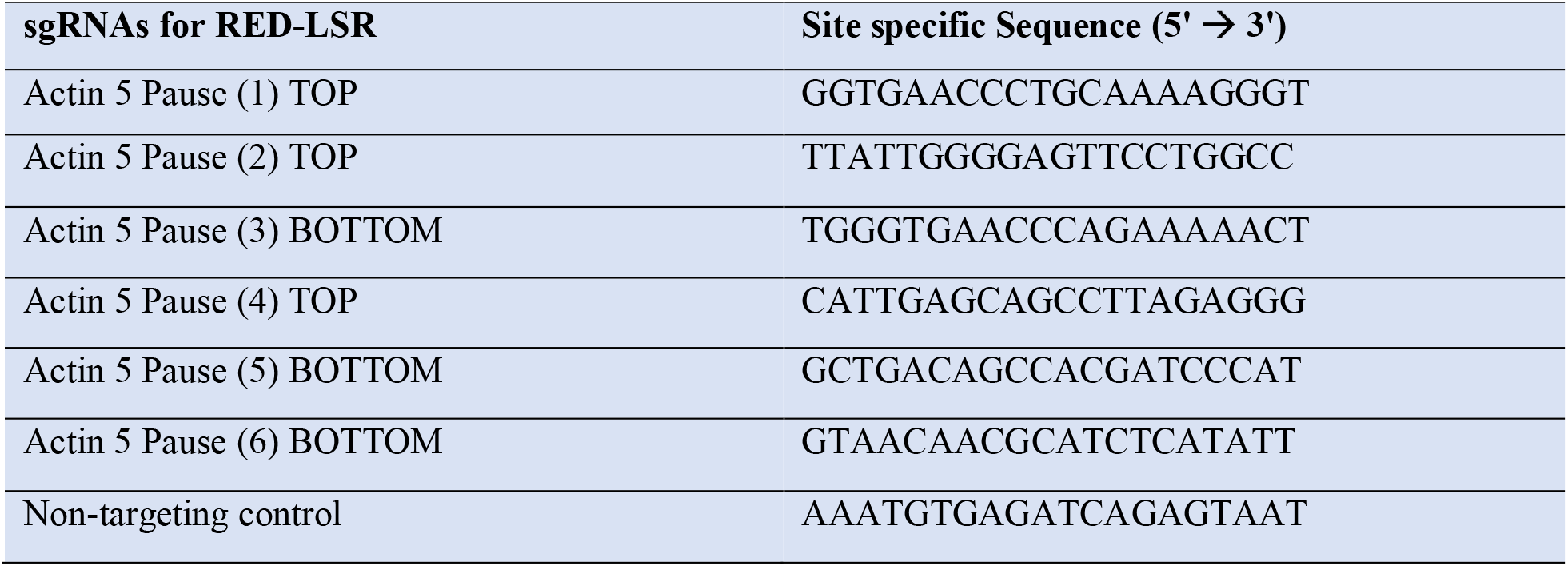
The sgRNA sequences used.

### Chromatin Immunoprecipitation (ChIP)

Chromatin immunoprecipitation (ChIP) of co-transfected HEK293T TREx™ cells was performed as previously described^3^. In brief, cells were crosslinked by adding 1% (*v/v*) formaldehyde at room temperature for 10 min, followed by 5 min incubation with 125 mM glycine. Cells were then lysed using lysis buffer (5mM PIPES, 85 mM KCl, 0.5% (*v/v*) NP-40, and complete protease-inhibitor cocktail). Cell pellets were resuspended in nuclear lysis buffer (50 mM Tris-HCl, 10 mM EDTA, 1% (*w/v*) SDS, and complete protease-inhibitor cocktail). Lysates were sonicated eight times for 20 s each at 40% amplitude at 4°C with intermittent 2 min incubations on ice. 10 μl of sheared chromatin from each sample was used as Input, while 50 ul was diluted 1/10 in immunoprecipitation dilution buffer (16.7 mM Tris-HCl pH 8.0, 0.01% (*w/v*) SDS, 167 mM NaCl, 1.2 mM EDTA, 1.1% (*v/v*) Triton X100, and complete protease inhibitor) and incubated with 5 μg of antibody on a rotator overnight at 4°C. Samples were then incubated at constant rotation with 25 μl of prewashed Dynabeads protein G (Life Technology, catalogue number 10004D) for 2 h at 4°C. Beads were washed once with low-salt wash buffer (20 mM Tris-HCl pH 8.0, 0.1% (w/v) SDS, 1% (v/v) Triton X-100, 2 mM EDTA pH 8.0, and 150 mM NaCl), once with high-salt wash buffer (20 mM Tris-HCl pH 8.0, 0.1% (w/v) SDS, 1% (v/v) Triton X-100, 2 mM EDTA pH 8.0, and 500 mM NaCl), once with LiCl wash buffer (10 mM Tris-HCl pH 8.0, 1% (v/v) NP-40, 1% (w/v) sodium deoxycholate, 1 mM EDTA pH 8.0, and 250 mM LiCl), and twice with TE buffer (10 mM Tris-HCl pH 8.0, and 1 mM EDTA pH 8.0) before two rounds of incubation with 100 μl of elution buffer for 15 min at room temperature. The eluates were incubated with 8 μl of 5 M NaCl on a rotator at 65°C overnight. We added 3 μl of 10 mg/ml RNaseA and incubated at room temperature 30 min, and then with 4 μl of 0.5 M EDTA, 8 μl of 1M Tris-HCl and 1 μl proteinase K (20 mg/mL) at 45°C for 2 h. DNA was purified using gel/PCR DNA fragment extraction (Geneaid, catalogue number DF300) and diluted with TE buffer. Samples were run with primers listed in Supplemental Information. ChIP-qPCR analysis was done as previously described^3^.

### DNA-RNA hybrid Immunoprecipitation (DRIP)

DNA-RNA hybrid immunoprecipitation (DRIP) of co-transfected HEK293T TREx™ cells was performed as previously described^3^. In brief, cells were washed with ice-cold PBS and centrifuged at 253 x *g* for 5 min. Cell pellets were resuspended and incubated in 1.6 ml of TE buffer, 0.05% SDS, and 5 μl of 20 mg/ml proteinase K overnight at 37°C. The genomic DNA was then purified using two rounds of phenol-chloroform and precipitated by adding 1/10 volume of 3 M NaOAc pH 5.2 and 2.4 volumes of 100% ethanol. The DNA fibre was washed with 70% ethanol, dried, resuspended in TE buffer, and then incubated with restriction enzyme mix (*Hin*dIII (NEB, R01045), *Eco*RI (ThermoFisher, ERO271), *Bsr*GI (NEB, R05755), *Xba*I (NEB, R01455), *Ssp*I (NEB, R0132), spermidine (Bioshop, catalogue number SPR070) and NEB buffer 2.1) overnight at 37°C. The digested genomic DNA was then purified using 3 M NaOAc pH5.2 and two rounds of phenol-chloroform, and precipitated by incubating with 2.4 volumes of cold 100% ethanol at - 20°C for 15 min, and centrifuged at maximum speed for 30 min at 4°C. The DNA pellet was washed with 70% ethanol and centrifuged at maximum speed for 5 min at 4°C. The dried DNA was resuspended in TE buffer, and 4.4 ug of the DNA was incubated with 350 ul TE buffer, 50 μl of 10x binding buffer (100 mM NaPO4 pH 7.0, 1.4 M NaCl, 0.5% (*v/v*) TritonX-100) and 10 μg of either mouse IgG or S9.6 antibody at 4°C overnight. Immunoprecipitation samples were incubated with previously washed Dynabeads for 2 h at 4°C. Samples were then washed three times with 1x binding buffer and eluted off the beads by incubation with DRIP elution buffer (50 mM Tris-HCl pH 8.0, 10 mM EDTA, 0.5% (*w/v*) SDS) and proteinase K for 45 min at 55°C. The DNA was then purified using gel/PCR DNA fragment extraction (Geneaid, catalogue number DF300) and qPCR of purified DNA was performed.

### RNA extraction

Co-transfected HEK293T TREx™ cells were washed with RNase-free PBS before RNA isolation using a Qiagen RNeasy mini Kit (Catalogue number 74104).

### Reverse transcription

Extracted RNA was reverse transcribed as previously described^3^. In brief, 1 μg of total RNA was treated with 1 μl of 10x DNase-I reaction buffer and 1 μl of DNase I Amp grade (1 U/μl; ThermoFisher, catalogue number 18068015), and then incubated for 15 min at room temperature. 1 μl of 25 mM EDTA was used to quench the reaction at 10 min for 65°C. A 10 μl reverse transcription reaction was then carried out using 10 mM deoxynucleoside triphosphate (dNTPs), 50 μM random nonamers (Sigma, catalogue number R7647), 500 ng total RNA, 5X first-strand buffer, 100 mM dithiothreitol (DTT), 40 U/μl RNaseOUT (Invitrogen, catalogue number 10777019), and 200 U/μl M-MLV reverse transcriptase (Invitrogen, catalogue number 28025013) at 25°C for 10 min, 37°C for 60 min and 70°C for 15 min.

### Quantitative PCR (qPCR)

Quantitative real-time PCR was performed using a Bio-Rad CFX connected Real-Time as previously described^3^. In brief, a 10 μl qPCR reaction is set up using SensiFAST SYBR No-ROX kit (FroggaBio/Bioline, catalogue number BIO-98050), forward and reverse primers (Table 2), and diluted DNA sample. PCR comprised of one cycle of 90°C for 5 min and 60°C for 30 s, followed by 39 cycles of 95°C for 5 s and 60°C for 30 s, and a final melt curve of 65°C to 95°C in 0.5°C steps at 5 s per step.

**Table 2.**
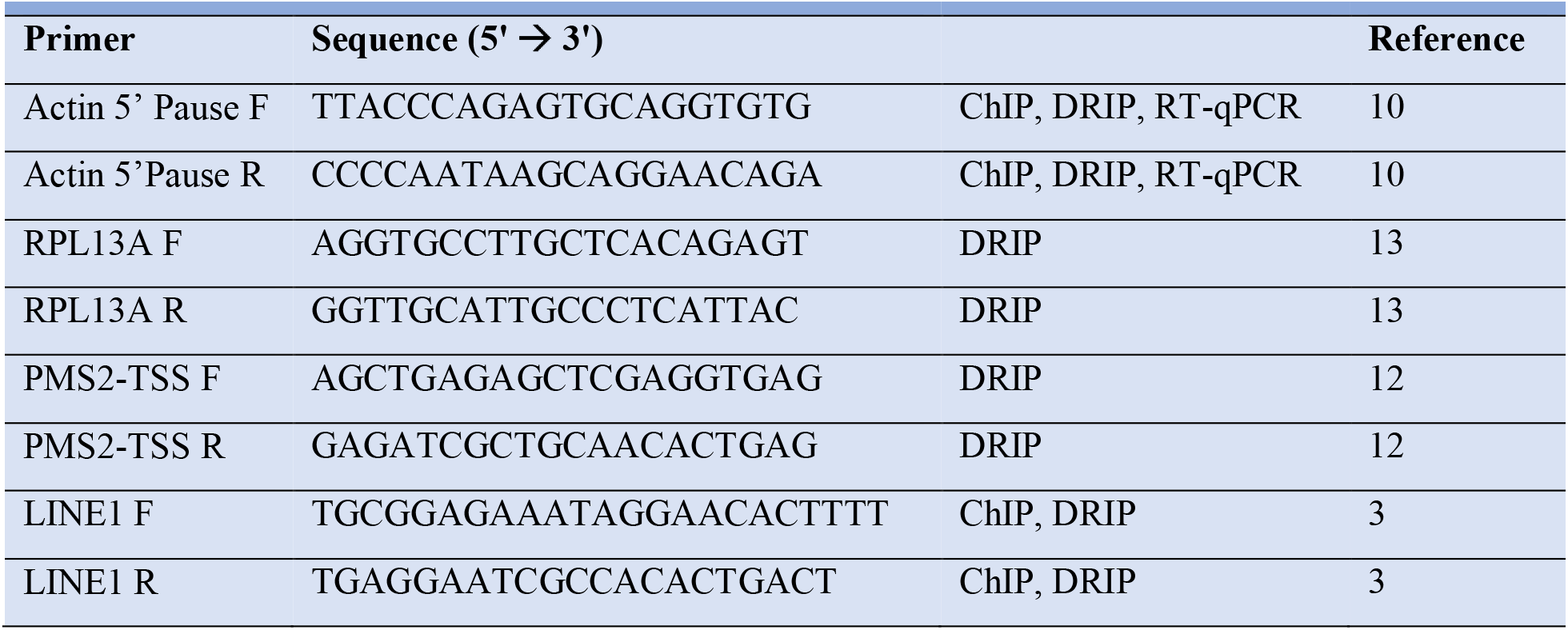
Primer sequences used.

## Data and materials availability

The RED/dRED-expressing plasmids used in this protocol have been deposited at Addgene (RED ID: 139835; dRED ID: 139836).

## Acknowledgments

NK and KJA were each supported by a Canadian Institutes of Health Research (CIHR) Vanier Doctoral Scholarship. NK was also partly supported by a CIHR CGS and Ontario Graduate Scholarship (OGS) awards. KJA was also supported by the Ruggles Innovation Award and Adel S. Sedra Award. This work was mainly supported by grants to KM from the CIHR (388041, 399687), Canada Research Chairs Program (CRC; 950-230661), and the Ontario Ministry of Research and Innovation (MRI-ERA; ER13-09-111).

## Author contributions

N.K. and KM conceived the paper. NK and KM wrote, and JNYC and KJA edited the manuscript. NK and JNYC performed experiments, and NK analyzed all data. KM supervised the research.

## Additional information

**Correspondence and requests for materials** should be addressed to KM.

